# Atypical Functional Connectivity in Tourette Syndrome Differs Between Children and Adults

**DOI:** 10.1101/459560

**Authors:** Ashley N. Nielsen, Caterina Gratton, Jessica A. Church, Nico U.F. Dosenbach, Kevin J. Black, Steven E. Petersen, Bradley L. Schlaggar, Deanna J. Greene

**Author notes:** Correspondence should be addressed to: Ashley N. Nielsen East Building Neuroimaging Laboratories 4525 Scott Ave, Suite 2220 St. Louis, MO 63110.

## Abstract

**Background:** Tourette syndrome (TS) is a neuropsychiatric disorder characterized by motor and vocal tics that typically change over development. Whether and how brain function in TS also differs across development has been largely understudied. Here, we used functional connectivity MRI to examine whole brain functional networks in children and adults with TS.

**Methods:** Multivariate classification methods were used to find patterns among functional connections that distinguish TS from controls separately for children and adults (total N = 202). We tested whether the patterns of connections that classify diagnosis in one age group (e.g., children) could classify diagnosis in another age group (e.g., adults). We also tested whether the developmental trajectory of these connections were altered in TS.

**Results:** Patterns of functional connections that distinguished TS from controls were generalizable to an age-matched independent test set, but not to other age groups. While diagnostic classification was successful in children and adults separately, the connections that best distinguished TS from controls were age-specific. When contextualized with typical development, some functional connections exhibited accelerated maturation in childhood TS, while others exhibited delayed maturation in adulthood TS.

**Conclusions:** Our results demonstrate that brain networks are differentially altered in children and adults with TS, and that the developmental trajectory of affected connections is disrupted. These findings further our understanding of neurodevelopmental trajectories in TS and carry implications for future applications aimed at predicting the clinical course of TS in individuals over development.

## INTRODUCTION

Tourette syndrome (TS) is a developmental neuropsychiatric disorder characterized by motor and vocal tics (1) that affects 1-3% of children (2–4). Tics are brief, unwanted, repetitive movements or noises that can be intrusive in daily life. On average, tic onset occurs at age 5–7 years, with tic severity peaking during late childhood/early adolescence (10-12 years). Tics usually continue into adulthood (5, 6), but with marked improvement or even remission after adolescence (7–11). However, symptom progression varies substantially across individuals, with a sizeable subgroup of patients (~60%) experiencing moderate to severe tics that persist into adulthood (9, 12). Understanding how the brain changes over the course of development in TS may provide insight into its clinical manifestation across development and aid prediction of the disorder’s trajectory in individuals.

Most neuroimaging studies of TS treat it as a singular disorder, unchanging across development, by grouping together patients from a wide age range (13–17) or focusing on a single age cohort (18–22), often by necessity. However, there is evidence that differences in brain structure and function in TS vary by age (23–25). Comparing the brain differences observed in children and adults with TS is necessary to reveal effects that are present in both age groups (i.e., “age-invariant” TS effects) as well as effects that differ between age groups (i.e., “age-specific” TS effects). Critically, a more complete understanding of the differences observed in children or adults with TS also requires taking into account typical maturational changes in the brain. Given a context of typical development, one can determine whether brain differences reflect atypically shifted development (e.g., accelerated or delayed maturation) or an anomalous difference not observed in typical development, potentially providing clues into etiology. While several TS neuroimaging studies have interpreted their findings in the context of brain maturity (23, 24, 26–28), few have included typical developmental comparisons to contextualize the differences observed in TS (29–31).

The potential presence of both maturity‐ and disorder-related differences in the brain in TS is made more complex by considering where these differences are localized. While many studies of TS have primarily identified differences within a select few brain regions or networks, the findings together suggest that TS involves many cortical and subcortical brain regions (for reviews, see (32, 33)). Thus, capturing the developmental trajectory of brain function in TS might be facilitated by a multivariate approach that combines information from many brain regions and identifies complex patterns in the data that distinguish individuals by diagnosis and/or age. Multivariate machine learning techniques have been applied to neuroimaging data in an attempt to identify patterns of diagnosis-related differences in neuropsychiatric disorders (34) and age-related differences in typical development (35–38). Notably, these methods require validation in an independent group of subjects to ensure that the identified differences do not represent idiosyncratic or spurious group differences (39), which is often not possible in small sample studies.

Here, we used a whole-brain, multivariate approach to investigate if and how brain networks in TS differ from controls in children and adults. Functional connectivity MRI, which measures the temporal correlations between spontaneous fluctuations in the blood oxygen level-dependent signals across the brain (40), was used to examine functional brain networks in separate cohorts of children and adults with TS. We previously demonstrated that multivariate approaches applied to functional connectivity can distinguish children with TS from controls (41) and typically developing children from adults (42, 43). In the present work, we use a similar approach, first validating that multivariate patterns of functional connections that distinguish TS and controls can generalize to an independent sample. Then, we test whether the patterns of functional connections that differ in TS in one age group (e.g., children) can also distinguish individuals with TS in the other age group (e.g., adults). Finally, we test whether the functional connections that differ in TS (in either children or adults) exhibit altered developmental trajectories by placing these differences in the context of typical development.

## MATERIALS AND METHODS PARTICIPANTS

### Participants

A total of 172 individuals with TS, ages 7.3-35.0 years, were recruited from the Washington University School of Medicine Movement Disorders Center and the Tourette Association of America Missouri chapter. After quality control assessments of the neuroimaging data (see below), 101 children, adolescents, and adults with TS were included (Table 1). A group of 101 control participants was selected from an extant database (*n*=487, ages 6.0–35.0 years, 206 males; recruited from the Washington University campus and surrounding community) and matched to the TS group on age, sex, IQ, handedness, and in-scanner movement (Table 1). Conditions commonly comorbid with TS (e.g., ADHD, OCD, anxiety) and medication use were not considered exclusionary for the TS group (44) (Table S1) but were for the control group. All participants completed assessments of IQ, and TS participants completed additional assessments of symptom severity for TS, ADHD, and OCD (Supplement 1.1). Adult participants and a parent or guardian for all child participants gave informed consent and all children assented to participation.

**Table 1.** Participant characteristics.

|  | **TS group** | **Control group** |
| --- | --- | --- |
| *N* | 101 | 101 |
| Male/Female | 62/39 | 61/40 |
| Age (Years) | 17.5 (7.6); 7.6-35.0 | 17.5 (7.5); 7.4-34.2 |
| Handedness (R/L) | 95/6 | 95/6 |
| IQ | 113 (13.3); 83-139 | 115 (13.8); 83-145 |
| Residual in-scanner movement (mean FD) | 0.11 (0.015); 0.063-0.14 | 0.11 (0.013); 0.067-0.13 |
| Amount of data ("good" frames) | 287.5 (100.8); 121-573 | 262.8 (102.1); 122-668 |
| YGTSS Total Tic Score | 17.4 (8.2); 0-37 | N/A |
| ADHD Rating Scale | 11.2 (10.2); 0-44 | N/A |
| CY-BOCS Score | 5.5 (6.2); 0-24 | N/A |
| Number on medications | 52 | 0 |
| Number with comorbidities | 67 | 0 |

Where applicable values are displayed as Average (Standard Deviation); Range
FD = Frame-wise Displacement (in millimeters) (45)
YGTSS = Yale Global Tic Severity Score (Total Tic Score) (46)
CY-BOCS Score = Children's Yale-Brown Obsessive-Compulsive Scale (47)

### Functional Connectivity Network Construction

Resting-state fMRI data were collected as participants viewed a centrally presented white crosshair on a black background. Participants were instructed to relax, look at the plus sign, and hold as still as possible. The duration and number of resting-state scans varied across participants (Supplement 1.2). Imaging data were collected using a 3T Siemens Trio Scanner with a 12-channel Head Matrix Coil. Images were pre-processed to reduce artifacts (48). Additional pre-processing steps were applied to the resting-state data to reduce spurious correlated variance unlikely related to neuronal activity. Stringent frame censoring (frame-wise displacement>0.2 mm) and nuisance regression (motion estimates, global signal, and individual ventricular and white matter signals) were used to reduce spurious individual or group differences in functional connectivity related to head movement in the scanner (49–51). Participants with at least 5 minutes of low-motion data were included. See Supplement 1.2–1.4 for details.

For each participant, resting-state time-courses were extracted from a set of 300 regions of interest (ROIs) (Figure 1) covering much of the cortex (52), subcortex, and cerebellum (available at https://greenelab.wustl.edu/datasoftware). Functional connectivity was measured as the correlation (Fisher z-transformed) between the resting-state time-courses for each pair of ROIs.

**Figure 1.**
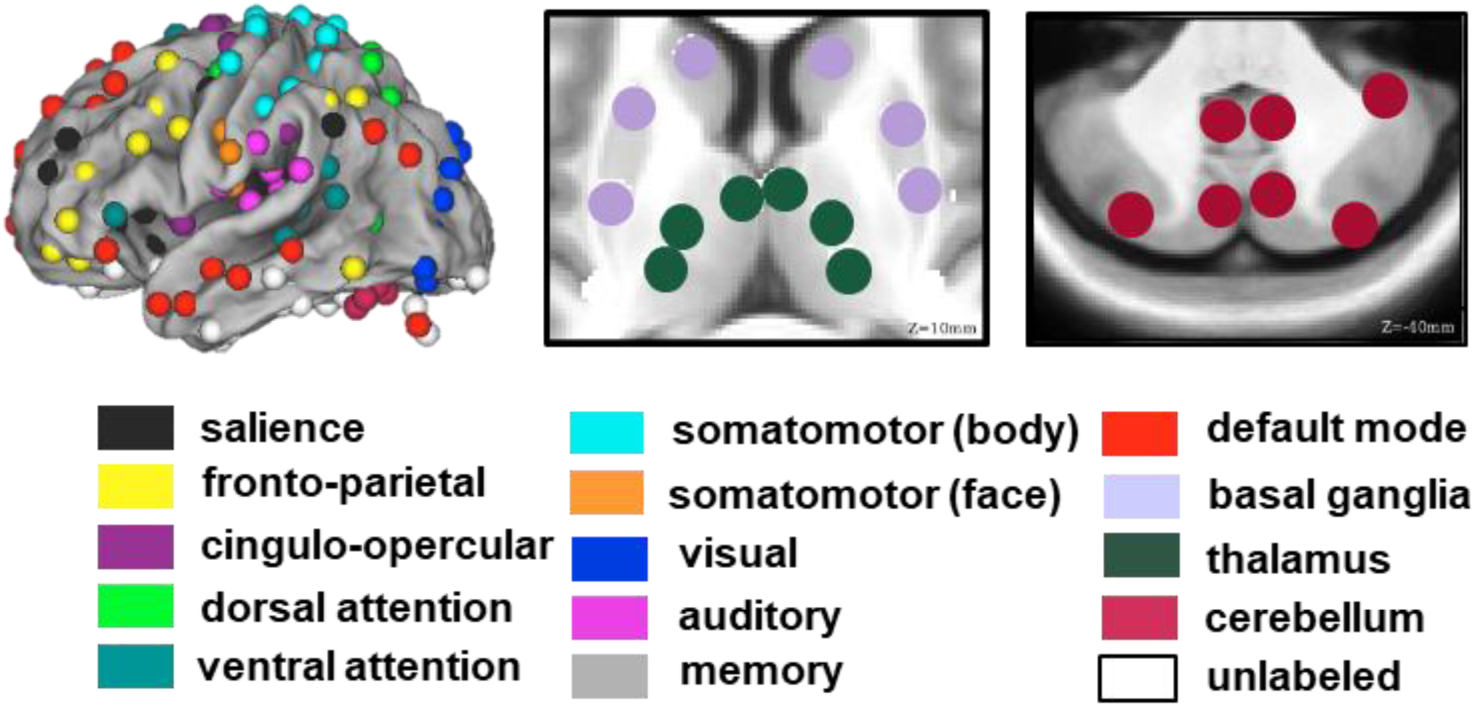
Regions of interest. Cortical regions were previously defined from a combination of task fMRI activation and resting-state fMRI studies (Power et al. 2011). Subcortical and cerebellar regions were defined from a combination of resting-state functional connectivity and review of the anatomical literature (Greene et al., 2014; Seitzman et al. (under review)). Cortical regions have been previously characterized as organizing into distinct functional networks (denoted by color).

### Support Vector Machine Learning

Support vector machine (SVM) learning was implemented (41–43) to distinguish individuals with TS from controls based on patterns of functional connections (Supplement 1.5). SVM classification is a powerful tool for finding differences across many features in a multivariate dataset (here, functional connections) that, in aggregate, best discriminate groups (here, TS vs. controls). Patterns of features that best distinguish individuals by group in a training set are weighted in the resulting classifier and can be subsequently applied to classify new test individuals. All 44,850 functional connections among the 300 ROIs were included as features.

Using SVM, three separate diagnostic classifiers were built to distinguish individuals with TS from controls using functional connectivity from three different training sets (Table 2). Leave-one-out cross validation (LOOCV) was used to assess classification accuracy within the training sets. The classifiers were then tested using independent test samples to answer several questions. A “YOUTH” diagnostic classifier was used to validate that patterns of functional connectivity that classify TS diagnosis are generalizable to an age-matched independent test set (see below). “CHILD” and “ADULT” diagnostic classifiers were used to test whether patterns of functional connectivity that classify TS diagnosis are age-specific or age-invariant (see below).

**Table 2.**
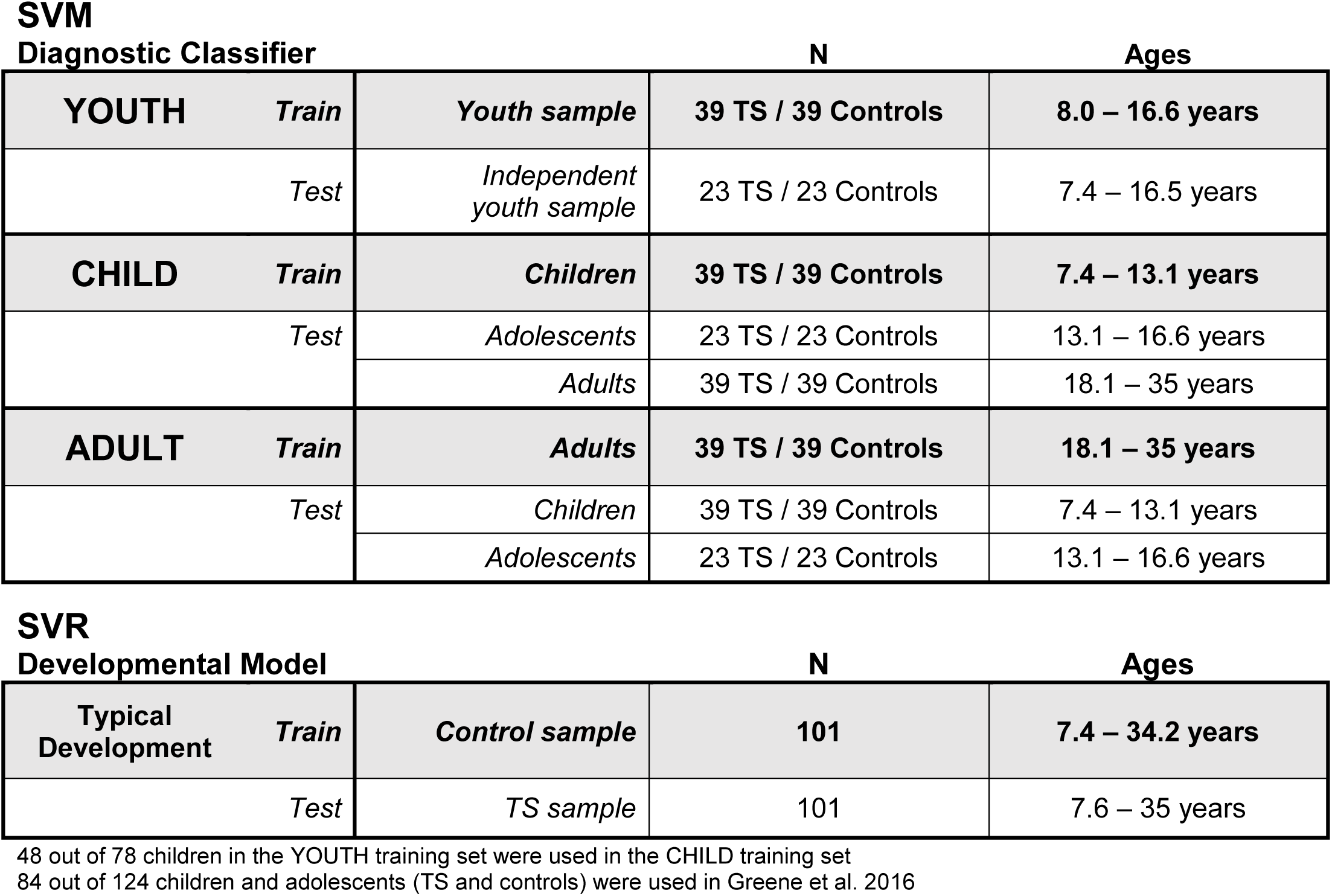
Overview of participants included in the training and testing sets for each diagnostic classifier and developmental model.

SVM classification can also be extended to find patterns among features that predict a continuous variable (here, age) with support vector regression (SVR). Using SVR, developmental models were built to predict age using functional connectivity from the controls (Table 2), and assessed with LOOCV, to generate a context of typical development.

### Validating diagnostic classification in an age-matched independent test set

We previously demonstrated that SVM can be used to classify children and adolescents with TS vs. controls based on patterns of functional connectivity (41). Here, we first wanted to ensure that the identified differences in functional connectivity characterize the disorder rather than idiosyncratic or spurious group differences within a specific sample. Because the TS sample contained many young individuals within an age range similar to that in Greene *et al.* 2016, we built a “YOUTH” diagnostic classifier trained to discriminate 39 children and adolescents with TS from 39 matched controls (8.0-16.6 years; Table 2) using SVM. The remaining 46 individuals were kept separate as an age-matched, independent youth sample (7.4-16.5 years; Table 2). We tested whether the YOUTH diagnostic classifier could accurately classify TS and controls in the independent youth sample.

### Testing for age-invariant or age-specific differences in functional connectivity in TS

We tested whether the patterns of functional connections that distinguished TS and controls were common or distinct between children and adults. Separate SVMs were used to build a CHILD diagnostic classifier trained to separate 39 children with TS from 39 matched controls (7.0-12.9 years; Table 2) and an ADULT diagnostic classifier trained to separate 39 adults with TS from 39 matched controls (18.0-35.0 years; Table 2). The remaining 23 adolescents with TS and 23 matched controls were kept as a separate adolescent test set (13.1-16.6 years; Table 2) to test whether the patterns of functional connections that classify diagnosis in children or adults can also classify diagnosis in adolescents.

We tested if the patterns of functional connections that distinguish TS from controls in one age group (child or adult) could generalize to accurately classify individuals in another age group. We evaluated whether the performance of a diagnostic classifier significantly differed across age groups using a binomial significance test (Supplement 1.6). As sex, comorbidities, and current medications were not matched across age groups (Table S2), we also tested whether the generalizability of the CHILD or ADULT diagnostic classifiers (or lack thereof) was driven by these characteristics (Supplement 2.1).

If the patterns of functional connections used to distinguish individuals with TS from controls in one age group are “age-specific,” the classifier should not generalize well to the other age group (i.e., the CHILD diagnostic classifier will not accurately distinguish adults with TS from adult controls, and vice versa). If these patterns are “age-invariant,” the classifier should generalize well to the other age group. We also directly tested for age-invariant differences using an ALL-AGES diagnostic classifier (Supplement 2.2).

We extracted the top 1000 (out of 44,850) most strongly weighted functional connections in each of the CHILD and ADULT diagnostic classifiers and examined the percentage overlap of those functional connections. Few overlapping connections would suggest age-specific differences between TS and controls, while many overlapping connections would suggest age-invariant differences (Supplement 2.3).

### Testing for anomalous or atypically shifted development of functional connectivity in TS

As previously reported, many functional connections vary systematically according to age in typical development (43). The functional connections that differ by diagnosis (TS vs. controls) may also vary according to age in typical development. To test this, we used SVR to build a developmental model using the top 1000 most strongly weighted functional connections from either the CHILD or ADULT diagnostic classifier, and tested if those features could also distinguish individuals by age in the control sample (7.4-34.2 years; Table 2). The developmental models built using the CHILD TS or ADULT TS features were also compared with developmental models built using randomly selected sets of functional connections to evaluate the utility of these specific features against a null model (Supplement 2.4).

We tested whether the developmental models built to predict age in the controls could also accurately predict age in the TS sample (7.6-35.0 years; Table 2). To benchmark the generalizability of age prediction to the TS sample, we also tested whether additional developmental models built to predict age in controls could accurately predict age in TS using 1) all 44,850 functional connections or 2) the top 1000 functional connections that differed most between control children and control adults (Supplement 2.5).

Determining if the most strongly weighted functional connections used for diagnostic classification can also predict age places the TS vs. control differences in the context of typical development, allowing interpretations pertaining to brain maturity. If the patterns of functional connections that distinguish TS from controls reflect an anomalous divergence unrelated to development, 1) the CHILD TS or ADULT TS features will not successfully predict age in controls or 2) those functional connections will predict age equivalently in both the control and TS samples. By contrast, if the patterns of functional connectivity that distinguish TS from controls reflect an atypically shifted developmental trajectory, the CHILD TS or ADULT TS features will predict age well in controls but inaccurately in TS. Predicted ages in TS that are older than in age-matched controls would indicate accelerated maturation of brain networks, while predicted ages in TS that are younger than in age-matched controls would indicate delayed or incomplete maturation of brain networks. Alternatively, if predicted ages in TS fluctuate near the mean age, the maturational changes present in typical development may be absent in TS.

## RESULTS

### Classification of TS vs. controls based on functional connectivity generalizes to an age-matched independent test set

Using SVM, we successfully classified individuals as TS or controls based on patterns of functional connectivity. The YOUTH diagnostic classifier, which included children and adolescents (8.0-16.6 years; Table 2), was 64% accurate when estimated with LOOCV, significantly above chance (*p*=0.01). Importantly, this diagnostic classifier successfully generalized to an independent youth sample of age-matched children and adolescents with 67% accuracy (Figure 2A). By demonstrating generalizability in an age-matched independent test set, we can better interpret the generalizability of the CHILD and ADULT diagnostic classifiers to different age groups; poor generalizability can likely be attributed to age-related differences in how brain networks are altered in TS rather than idiosyncratic group differences related to data quality or overfitting.

**Figure 2.**
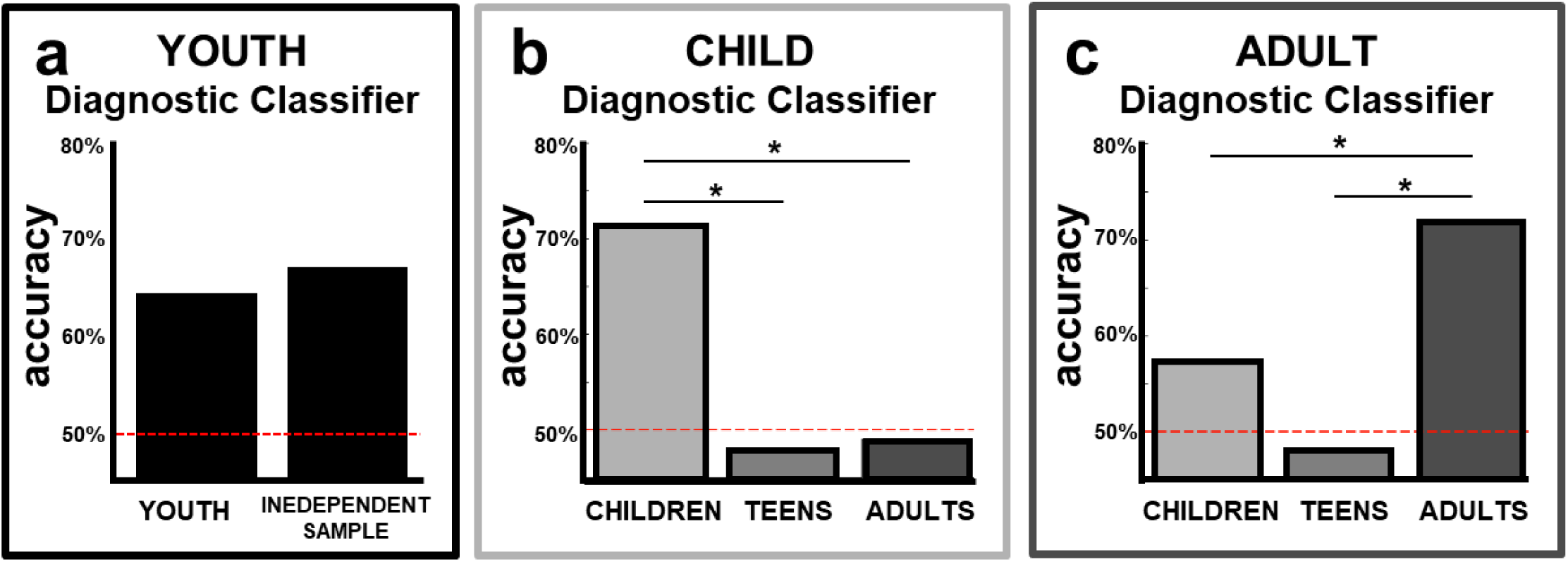
Functional connections that best distinguished TS from controls were age-specific. **a**.) Performance of the YOUTH diagnostic classifier was significantly better than chance in the independent sample (*p* = 0.01). **b**.) Performance of the CHILD diagnostic classifier was not significantly better than chance in adolescents (accuracy: 48%, sensitivity: 91%, specificity: 4%, *p* = 0.48) or adults (accuracy: 49%, sensitivity: 97%, specificity: 0%, *p* = 0.43) and was significantly less accurate in classifying adolescents and adults than children (adolescents: *p* < 0.001 ; adults: *p* < 0.001). c.) Performance of the ADULT diagnostic classifier was not significantly better than chance in adolescents (accuracy: 48%, sensitivity: 17%, specificity: 78%, *p* = 0.49) or children (accuracy: 57%, sensitivity: 31%, specificity: 85%, *p* = 0.11) and was significantly less accurate in classifying children and adolescents than adults (adolescents: *p* < 0.001 ; children: *p* = 0.012).

### Patterns of functional connections can classify TS diagnosis in children and in adults, but do not generalize across age groups

The CHILD diagnostic classifier (7.4-13.1 years; Table 2) was 71% accurate (LOOCV, *p*<0.001). The ADULT diagnostic classifier (18.1-35.0 years; Table 2) was 72% accurate (LOOCV, *p*<0.001). However, neither classifier accurately classified TS diagnosis in the other age groups (Figure 2B-C). Specifically, the CHILD diagnostic classifier did not distinguish TS from controls in adolescents (accuracy: 48%, *p*=0.48) or adults (accuracy: 49%, *p*=0.43). Similarly, the ADULT diagnostic classifier did not distinguish TS from controls in adolescents (accuracy: 48%, *p*=0.49), though it was slightly better in children (accuracy: 57%, *p*=0.11). Classification of the other age groups was significantly less accurate than classification in the training sample (see Figure 2). Given the successful generalizability of the YOUTH diagnostic classifier (described above), poor generalizability is likely not solely related to data quality or overfitting. Moreover, poor generalizability was not driven by sex, comorbid disorders, or medication status (Supplement 2.1, Table S3). These results suggest that the CHILD and ADULT diagnostic classifiers relied on age-specific differences in functional connectivity to best discriminate TS from controls. We also found evidence for age-invariant differences in functional connectivity in TS (Supplement 2.2). However, those age-invariant patterns were not the primary features used to distinguish TS and controls when considering children and adults separately.

### Top functional connections that distinguish TS and controls were distinct in children and adults

Regions associated with the top weighted functional connections from the CHILD and ADULT diagnostic classifiers are displayed in Figure 3, and show that these functional connections were within and between many different functional networks (Supplement 2.3, Figure S3). Only 33 (3%) of the top 1000 functional connections overlapped between the CHILD and ADULT diagnostic classifiers (Figure 3C), indicating different patterns of region involvement (Figure 3A-B) and providing further evidence that the functional connections involved in TS differ in children and adults.

**Figure 3.**
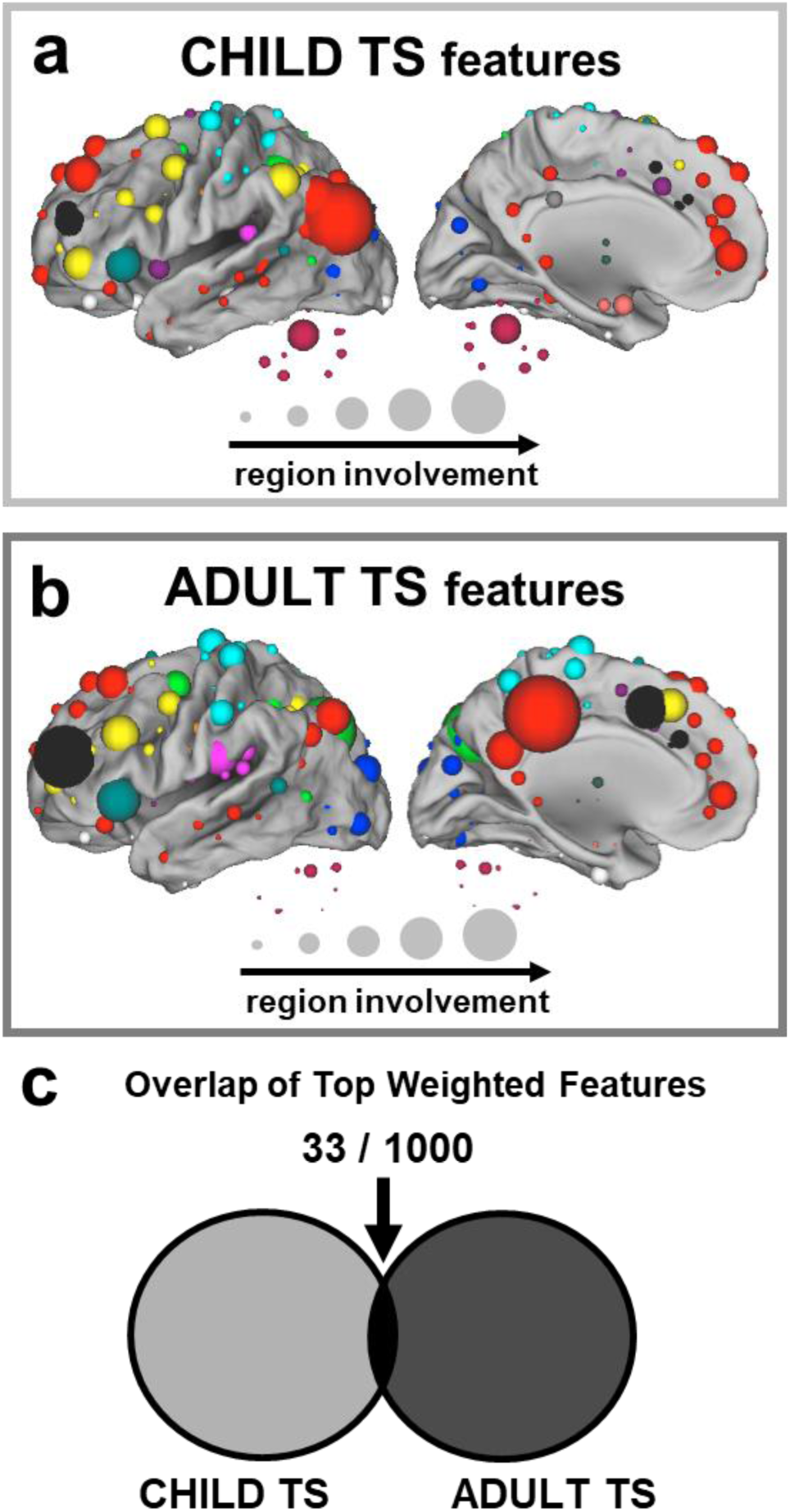
Functional connections that best distinguished TS from controls differed between children and adults. **a**.) Regions are shown from the top weighted 1000 functional connections used to distinguish TS from controls in the CHILD diagnostic classifier. The size of each sphere represents region involvement (i.e., number of functional connections in the feature set involving a region). Region colors indicate the network to which that region belongs, labeled in Figure 1. **b**.) Regions are shown from the top weighted 1000 functional connections used to distinguish TS from controls in the ADULT diagnostic classifier. The size of each sphere represents region involvement and the color represents network affiliation. **c**.) The overlap of the top weighted functional connections from the CHILD and ADULT diagnostic classifiers was only 33 out of 1000.

### Functional connections that differ in TS reflect atypically shifted development

Using SVR, the top weighted functional connections from the CHILD diagnostic classifier and the ADULT diagnostic classifier were each able to predict age well in the controls (CHILD: *r*=0.62, *R*^2^=0.39, *p*<0.001; ADULT: *r*=0.74, *R*^2^=0.55, *p*<0.001; Figure 4, *red*) when evaluated against a null model (Supplement 2.4, Figure S4). By contrast, these developmental models did not predict age well in TS. Specifically, the developmental model built to predict age in controls using the CHILD TS features did not significantly predict age in TS, r=0.11, *R*^2^=0.012, *p*=0.27 (Figure 4A, *blue*) such that the children with TS were inaccurately predicted as older than age-matched controls. Note that these predicted ages were shifted above the age expected if predicted spuriously (Supplement 2.6, Figure S5), suggesting accelerated maturation of these functional connections in childhood TS. The developmental model built to predict age in controls using the ADULT TS features also did not significantly predict age in TS, *r*=0.11, *R*^2^=0.013, *p*=0.27 (Figure 4B, *blue*) such that the adults with TS were inaccurately predicted as younger than age-matched controls. These predicted ages were shifted below the mean age (Supplement 2.6, Figure S5), suggesting delayed maturation of these functional connections in adulthood TS.

**Figure 4.**
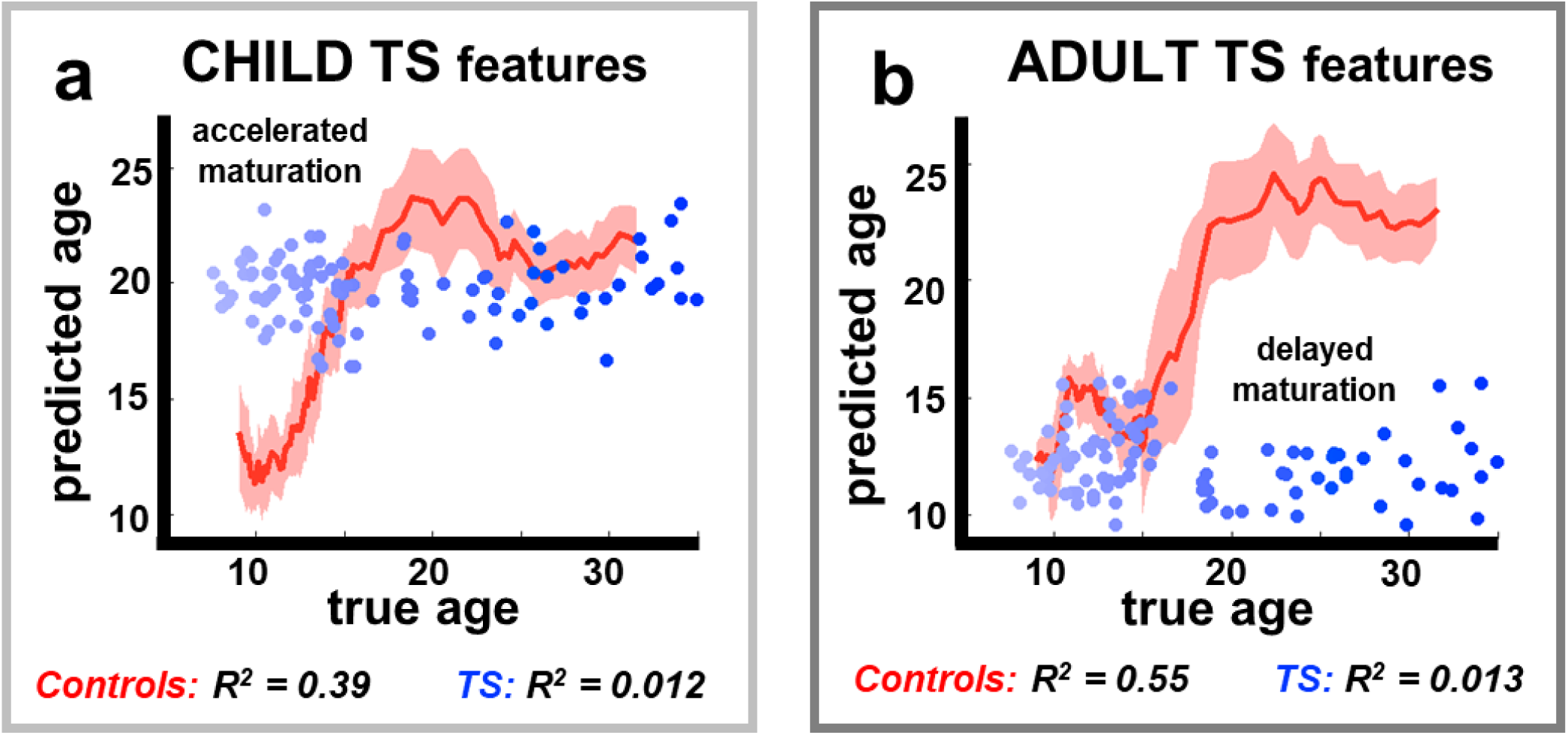
Functional connections that best distinguished TS from controls reflect atypically shifted development. **a**.) The developmental model built using CHILD TS features was able to predict age well in the control sample *(red)* but not in the TS sample (*blue*). Predicted ages of children with TS were older than the predicted ages of age-matched controls indicating accelerated maturation of the CHILD TS features. **b**.) The developmental model built using ADULT TS features was able to predict age well in the control sample (*red*) but not in the TS sample (*blue*). Predicted ages of adults with TS were younger than the predicted ages of age-matched controls indicating delayed or incomplete maturation of the ADULT TS features.

Not all development of functional connectivity was disrupted in TS. We found that additional developmental models could accurately predict age in the TS sample using 1) whole-brain functional connectivity (*r*=0.71, *R*^2^=0.50, *p*<0.001) and 2) the functional connections that differ most between control children and adults (*r*=0.62, *R*^2^=0.38, *p*<0.001; Supplement 2.5). Thus, only the top functional connections used to distinguish TS from controls in each age group demonstrated altered developmental trajectories in TS.

## DISCUSSION

In the present work, we applied multivariate machine learning methods to resting-state functional connectivity MRI data to understand how functional brain organization is altered in TS over development. We found that the patterns of functional connections that best distinguished TS from controls were generalizable to an age-matched independent sample, but not to other age groups. Rather, the functional connections involved in TS differed between children and adults, suggesting they are age-specific. In addition, we found that these functional connections reflected atypical development in TS. Specifically, those functional connections that differed the most in childhood TS exhibited accelerated maturation (i.e., resembled brain networks of older subjects), while those that differed the most in adulthood TS exhibited delayed maturation (i.e., resembled brain networks of younger subjects). By directly examining TS across a wide age range (7-35 years), comparing children to adults, and contextualizing these results with typical development, our findings provide evidence that the neural underpinnings of TS differ in childhood and adulthood, and involve changes to the typical brain maturation timeline.

It has been argued that childhood and adulthood TS are fundamentally different, given the common clinical trajectory in which many patients experience significant improvement or remission in adulthood (53). Our results extend this argument to the brain’s functional connections. Past studies have also identified age-specific effects in TS, yet primarily within single brain regions. For example, some cortical regions (dorsal prefrontal, orbitofrontal, parieto-occipital cortex) exhibit distinct, even sometimes opposing, volumetric differences in children and adults with TS (26). Previous research has also shown that motor excitability is selectively altered in children with TS (23) and atypical development of fronto-striatal self-regulatory signals only emerges in adulthood TS (24). These findings in combination with ours suggest that treatments may need to be tailored differently for children and adults with TS.

We also characterized functional connectivity in TS in the context of typical development. In childhood TS, we found differences indicative of accelerated development. It has been proposed that living with chronic tics accelerates the maturation of control systems in children with TS as a result of the need to regularly suppress tics (53, 54). In line with this idea, previous studies have reported enhanced cognitive control as well as putatively adaptive changes in brain function and structure in children with TS (55–57). In heathy children, cognitive training yields modifications of the intrinsic connectivity among brain networks (58). Thus, it is possible that the development of compensatory tic-suppression mechanisms is reflected in the patterns of functional connectivity that best distinguish children with and without TS. It is possible that these alterations support the improvement of tic symptoms during adolescence and early adulthood experienced by many patients (59).

In adulthood TS, we found differences in functional connectivity indicative of delayed maturation. Thus, adults that experience persistent tics may have maladaptive brain function that either developed with prolonged symptoms or led to the prolonged symptoms. As mentioned above, some argue that childhood TS and adulthood TS are fundamentally different, given the commonly held belief that most patients with TS experience substantial symptom improvement or remission into adulthood (9). Therefore, by studying a sample of adults with current tics, we may have captured the subsample who do not experience remission. By contrast, any sample of children with TS will include a mixture of individuals whose tic symptoms will go on to improve and those whose tics will persist. However, there is evidence that remission is likely much rarer than previously estimated (10%, rather than 40%; (12)), and in our sample, many of the adults with TS reported improvement from childhood even if they did not report remission. Longitudinal data and studies of adults with remitted tics are necessary to determine whether immature brain function in adulthood TS is a cause or consequence of prolonged symptom burden. There have been previous reports of immature brain structure and function in TS (28, 30, 60, 61). However, methodological concerns related to head motion artifact in MRI data have called some of these conclusions into question (49, 62–65). In the present study, we implemented strict processing methods that have been shown to best mitigate the artifactual effects of motion (49, 51). Evidence for altered maturation of functional connectivity in TS remains even when potential artifactual confounds have been addressed.

Notably, not all maturation of functional connectivity was altered. When the complete set of functional connections across the brain were included in the developmental model, age was predicted well in both TS and controls. Further, those functional connections that varied with age the most in controls could also predict age well in TS. Thus, only specific patterns of functional connections – those that best discriminated TS and controls within each age group – exhibited shifted developmental trajectories in TS, while much of the typical maturation of functional connectivity was preserved. This finding may correspond to the clinical observation that although TS can involve diminished academic achievement and quality of life, most individuals with TS lead relatively normal lives (66, 67).

It is important to note that our TS sample was heterogeneous with respect to comorbid neuropsychiatric disorders and medication status, representative of the TS population (44, 68). As brain network function can be affected by medications (69) and other neuropsychiatric conditions (70), the diagnostic classifiers here might have included medication-induced or comorbidity-related differences in brain function between the TS and control groups. Additionally, our child and adult samples differed with respect to sex; the children included more boys than girls, while the adults were more balanced. This difference reflects epidemiological data, as the sex imbalance (4:1 male:female) reported in childhood TS is attenuated in adulthood TS (71). Nevertheless, examination of the misclassified individuals demonstrated that poor generalizability across age groups was not driven by these factors. Future studies with larger samples will be useful for directly parsing the influence of medications, comorbidities, and sex on brain function in TS.

The success of multivariate machine learning classification applied to functional brain networks holds promise for clinical application of these methods. Given the heterogeneity in the developmental course of TS symptoms, there is a great need to predict future clinical outcome for individuals. Being able to predict whether a given child with tics will go on to improve or not would have high clinical utility, providing important information to families, guiding treatment plans, and affording the opportunity for early intervention. Our findings suggest that functional connectivity contains signals that can be used for these types of predictions, and that the best predictions will likely rely upon modeling these effects in a rich typical developmental context.

## ACKNOWLEDGEMENTS

We thank Rebecca Coalson, Rebecca Lepore, Kelly McVey, Jonathan Koller, Annie Nguyen, Catherine Hoyt, Lindsey McIntyre, and Emily Bihun for assistance with data collection, the children and adults who participated in this study and their families, and Deanna Barch for her input on this manuscript. This project was supported by: Tourette Association of America fellowships (DJG, JAC), Tourette Association of America Neuroimaging Consortium pilot grant (KJB, BLS), Tourette Association of America research grant (DJG), NARSAD Young Investigator Award (DJG), NIH K01MH104592 (DJG), NIH R21MH091512 (BLS), NIH R01HD057076 (BLS), NIH R01NS046424 (SEP), NIH R21 NS091635 (BLS, KJB), NIH R01MH104030 (KJB, BLS), K12HD076224 (NUFD, Scholar of the Child Health Research Center at Washington University), NIH K23NS088590 (NUFD), NIH F32NS065649 (JAC), NIH F32NS092290 (CG), NIH F32NS656492, NIH K23DC006638, P50 MH071616, P60 DK020579-31 American Hearing Research Foundation, and The Simons Foundation Autism Research Initiative (“Brain circuitry in Simplex Autism,” SEP). Research reported in the publication was supported by the Eunice Kennedy Shriver National Institute of Child Health & Human Development of the National Institutes of Health under Award Number U54 HD087011 to the Intellectual and Developmental Disabilities Research Center at Washington University. The content is solely the responsibility of the authors and does not necessarily represent the official views of the National Institutes of Health.

## 1. SUPPLEMENTAL METHODS

### 1.1 Participants

A total of 101 children and adults with Tourette syndrome (TS) and 101 healthy control children and adults were included in the present study. All participants were native English speakers. All participants underwent a 2-scale brief assessment of IQ (WASI). For TS participants, the experimenter completed the following measures of “past week” symptom severity: Yale Global Tic Severity Score (Total Tic Score) (46), Children’s Yale-Brown Obsessive Compulsive Scale (47), and ADHD Rating Scale (72). All participants self‐ or parent-reported any history of neuropsychiatric diagnoses and current medications (Table S1). For the control participants, any history of neuropsychiatric or neurological diagnoses prohibited participation in the study.

**Table S1.** Comorbid diagnoses and current medications in participants with TS.

|  | **Children with TS**<br>*N* = 39; (7.4 – 13.1 years) | **Adolescents with TS**<br>*N* = 23; (13.1-16.6 years) | **Adults with TS**<br>*N* = 39; (18.0-35 years) |
| --- | --- | --- | --- |
| **Comorbid Diagnosis** |  |  |  |
| ADHD/ADD | 18 | 11 | 11 |
| OCD | 9 | 8 | 13 |
| Anxiety Disorders | 5 | 4 | 9 |
| Depression | 1 | 2 | 9 |
| ODD | 0 | 2 | 0 |
| Migraines | 1 | 0 | 6 |
| **Medications** |  |  |  |
| Centrally acting adrenergic agents | 13 | 9 | 3 |
| Stimulants | 10 | 3 | 7 |
| Anti-depressants | 2 | 5 | 6 |
| Anti-anxiety | 1 | 0 | 2 |
| Antipsychotics | 0 | 1 | 1 |

### 1.2 Imaging Acquisition

Data were acquired on a Siemens 3T Trio scanner (Erlanger, Germany) with a Siemens 12-channel Head Matrix Coil. Each child was fitted with a thermoplastic mask fastened to the head coil to help stabilize head position. T1-weighted sagittal MP-RAGE structural images in the same anatomical plane as the BOLD images were obtained to improve alignment to an atlas (1 sequence acquisition for each of the 101 control participants (child, adolescent, and adult) and for 88 of the TS participants (child, adolescent, adult): slice time echo = 3.06 ms, TR = 2.4 s, inversion time = 1 s, flip angle = 8°, 176 slices, 1 × 1 × 1 mm voxels; 2 sequence acquisitions for each of the 13 remaining child and adolescent TS participants: slice time echo = 2.34 ms, TR = 2.2 s, inversion time = 1 s, flip angle = 7°, 160 slices, 1 × 1 × 1 mm voxels). Functional images were acquired using a BOLD contrast-sensitive echo-planar sequence (TE = 27 ms, flip angle = 90°, in-plane resolution 4x4 mm; volume TR = 2.5 s). Whole-brain coverage was obtained with 32 contiguous interleaved 4 mm axial slices. Steady-state magnetization was assumed after 4 volumes. For most participants, 2-4 resting state scans lasting 5-5.5 min each were acquired, but the duration of each scan ranged from 3.2 minutes to 30 minutes. In the TS group, 388 ± 61.5 (range 264-528) total functional volumes were acquired, and in the control group, 372 ± 130 (range 260-724) total functional volumes were acquired.

### 1.3 Imaging preprocessing

Functional images from each participant were preprocessed to reduce artifacts (Shulman et al. 2010). These steps included: (i) temporal sinc interpolation of all slices to the temporal midpoint of the first slice, accounting for differences in the acquisition time of each individual slice, (ii) correction for head movement within and across runs, and (iii) intensity normalization of the functional data was computed for each individual via the MP-RAGE T1-weighted scans. Each run was then resampled in atlas space on an isotropic 3 mm grid combining movement correction and atlas transformation in a single interpolation. The target atlas was created from thirteen 7-9 year old children and twelve 21-30 year old adults using validated methods (Black et al. 2004). The atlas was constructed to conform to the Talairach atlas space.

### 1.4 Functional Connectivity Preprocessing

Several additional pre-processing steps were applied to reduce spurious variance unlikely to reflect neuronal activity (Fox et al. 2009). These functional connectivity pre-processing steps included: (i) demeaning and detrending each run, (ii) multiple regression of nuisance variables, (iii) frame censoring (discussed below) and interpolation of data within each run, (iv) temporal band-pass filtering (0.009 Hz < f < 0.08 Hz), and (v) spatial smoothing (6 mm full width at half maximum). Nuisance variables included motion regressors (e.g. original motion estimates, motion derivatives, and Volterra expansion of motion estimates), an average of the signal across the whole brain (global signal), individualized ventricular and white matter signals, and the derivatives of these signals.

We applied a procedure determined and validated to best reduce artifacts related to head motion (Power et al. 2014; Ciric et al. 2017). Specifically, frame-by-frame head displacement (FD) was calculated from preprocessing realignment estimates, and frames with FD > 0.2 mm were removed. An FD threshold of 0.2 mm was chosen because it best reduced the distance-dependence related to individual differences in head motion (mean FD) in this developmental dataset, as assessed using procedures from Power et al. (2012) and Ciric et al. (2017). Data were considered usable only in contiguous sets of at least 3 frames with FD < 0.2 and a minimum of 30 frames within a functional run. Motion-contaminated frames were censored from the continuous, processed resting-state time series before computing resting-state correlations. Notably, the global signal was included as a nuisance regressor (mentioned above) in order to further reduce global, motion-related spikes in BOLD data (Power et al. 2014; Ciric et al. 2017) and reduce patterns of spurious functional connectivity that might be utilized for prediction with machine learning (Nielsen et al. 2018).

### 1.5 Parameters for Support Vector Machine Learning

The parameters used for support vector machine (SVM) learning were the same as those used in Dosenbach et al. 2010 and Greene et al., 2016. We used the Spider Machine Learning Toolbox implemented in Matlab for SVM training and testing. In SVMs, each of the samples (here, participants) is treated as a point in multidimensional space defined by as many dimensions as features (here, 44,850 functional connections). In training an SVM classifier, a penalty is incurred for misclassified data in the training set (points on the wrong side of the multivariate decision boundary). The parameter C describes the margin used in training. For a larger C, a larger penalty is assigned to misclassification errors. All SVM classifications described in this work used soft-margin SVMs with the default setting of C = 1. Leave-one-out cross validation (LOOCV) was used to assess how well a classifier can distinguish individuals from different groups. In turn, each individual was removed from the training set, a diagnostic classifier was built to distinguish TS from controls in the remaining participants, and the left out subject was classified with the resulting diagnostic classifier.

We empirically tested whether a diagnostic classifier performed significantly above chance. We randomly sorted individuals with and without TS into two classes and trained a classifier distinguish the two arbitrary classes. LOOCV was used to determine the diagnostic classification accuracy of each classifier (expected accuracy is near 50%). We repeated this randomization process 100 times. By comparing the observed classification accuracy in the CHILD or ADULT diagnostic classifiers to the classification accuracy of the diagnostic classifiers trained with arbitrary classes, we can determine whether the CHILD or ADULT diagnostic classifier can discriminate TS from controls above chance.

The parameters used for support vector regression (SVR) were the same as those used in Dosenbach et al. 2010 and Nielsen et al. 2018. SVR retains some of the main features of binary SVM classification. In SVR, a penalty is incurred for data that is too far from the regression line in multivariate space. Epsilon-insensitive SVR defines a tube of width epsilon around the regression line in multivariate space. Any data points (i.e., subjects) within this tube carry a loss of zeros, meaning there is no penalty. In SVR, the C parameter controls the trade-off between how strongly subjects beyond the epsilon insensitive tube are penalized and the flatness of the regression line (larger C allows the regression line to be less flat). All SVR predictions described here used epsilon-insensitive SVRs with the Spider Machine Learning Toolbox default setting of C = Infinity and epsilon = 0.00001.

### 1.6 Binomial Significance Test

To determine whether the performance of a diagnostic classifier significantly differed from an expected performance, we used a binomial significance test. We determined the probability density function for an observed classification accuracy *x*, given *n* independent test subjects and *p*, the expected accuracy of a diagnostic classifier, as follows.

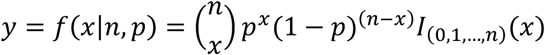

The result, *y*, is the probability of observing *x* in *n* independent trials, where the probability of correctly classifying TS in any given subject is *p*.

This approach was used to assess whether:

A. CHILD diagnostic classifier performed significantly differently in adolescents and adults than in children
B. ADULT diagnostic classifier performed significantly differently in children and adolescents than in adults
C. CHILD and ADULT diagnostic classifiers performed significantly differently than the ALL-AGES diagnostic classifier (see Supplement 2.2 *ALL-AGES Diagnostic Classifier*)
D. Misclassification of individuals according to sex, comorbid disorders, or current medications was significantly different than expected given the composition of the test set (see Supplement 2.1 *Misclassification of potentially confounding characteristics*).

## 2. SUPPLEMENTAL ANALYSES

### 2.1 Misclassification of potentially confounding characteristics

As shown in Figure 2, the CHILD and ADULT diagnostic classifiers did not generalize well to other age groups. Notably, these classifiers systematically misclassified individuals from other age groups, as evidenced by the imbalanced sensitivity and specificity. The CHILD diagnostic classifier misclassified control adolescents and adults as TS, while the ADULT diagnostic classifier misclassified children and adolescents with TS as controls. The CHILD diagnostic classifier misclassified 24 out of 46 adolescents (2 TS / 22 control) and 40 out of 78 adults (1 TS / 39 control). The ADULT diagnostic classifier misclassified 24 out of 46 adolescents (19 TS / 5 control) and 33 out of 78 adults (27 TS / 6 control). While the total TS and control samples were matched on age, sex, IQ, handedness, and in-scanner movement, not all of these characteristics were matched across age groups (Table S2). Sex ratio, frequency of comorbid disorders, and the number of individuals currently taking medications varied across the groups of children, adolescents, and adults.

**Table S2.** TS participant characteristics per age group.

|  | Age range | Sex Ratio | Comorbidities | Medications | YGTSS |
| --- | --- | --- | --- | --- | --- |
| **Children** | 7.4 – 13.1 years | 61 M: 17 F | 23 | 19 | 17.5 (8.1) |
| **Adolescents** | 13.1 – 16.6 years | 30 M: 16 F | 16 | 14 | 18.2 (8.2) |
| **Adults** | 18.1 – 35 years | 32 M: 46 F | 28 | 18 | 16.8 (8.4) |

To test whether these characteristics affected classification, we first used a binomial significance test to determine if the composition of misclassified individuals from the CHILD or ADULT diagnostic classifiers significantly differed from the overall composition of the test set. A summary of these results is shown in Table S3. Overall, misclassified individuals were representative of the test set containing largely the same sex ratio and percentage of individuals with comorbidities or current medications.

**Table S3.** Comparing the composition of misclassified individuals from the CHILD and ADULT Diagnostic Classifiers to that of the original test set.

| **CHILD**<br>**Diagnostic**<br>**Classifier** | **Test Set** | **Number**<br>**misclassified** | **Composition of**<br>**Test Set** | **Composition of**<br>**Misclassifications** | **Difference**<br>**uncorrected**<br>**p-value** |
| --- | --- | --- | --- | --- | --- |
| **Sex**<br>% males | *Adolescents* | 24 | 65% | 63% | 0.16 |
|  | *Adults* | 40 | 41% | 40% | 0.13 |
| **Comorbidities**<br>% with comorbidities | *TS Adolescents* | 2 | 70% | 50% | 0.42 |
|  | *TS Adults* | 1 | 72% | 100% | 0.72 |
| **Medications**<br>% on medications | *TS Adolescents* | 2 | 61% | 50% | 0.48 |
|  | *TS Adults* | 1 | 46% | 0% | 0.54 |
| **ADULT**<br>**Diagnostic**<br>**Classifier** | **Test Set** | **Number**<br>**misclassified** | **Composition of**<br>**Test Set** | **Composition of**<br>**Misclassifications** | **Difference**<br>**uncorrected**<br>**p-value** |
| **Sex**<br>% males | *Children* | 33 | 78% | 64% | 0.021* |
|  | *Adolescents* | 24 | 65% | 63% | 0.16 |
| **Comorbidities**<br>% with comorbidities | *TS Children* | 27 | 59% | 59% | 0.15 |
|  | *TS Adolescents* | 19 | 70% | 63% | 0.16 |
| **Medications**<br>% on medications | *TS Children* | 27 | 49% | 41% | 0.15 |
|  | *TS Adolescents* | 19 | 61% | 53% | 0.14 |

As the sex ratio of the misclassified children using the ADULT diagnostic classifier differed from the sex ratio of the entire set of children (*p* = 0.021, *uncorrected*), we wanted to ensure that poor generalizability was not due to sex differences between age groups. Thus, we built sex-matched CHILD and ADULT diagnostic classifiers and tested how well these diagnostic classifiers generalized to other age groups. The sex-matched training sets were smaller (n=66 rather than n=78) and sampled from a broader age range (Table S4) than the training sets used in the main text.

**Table S4.**
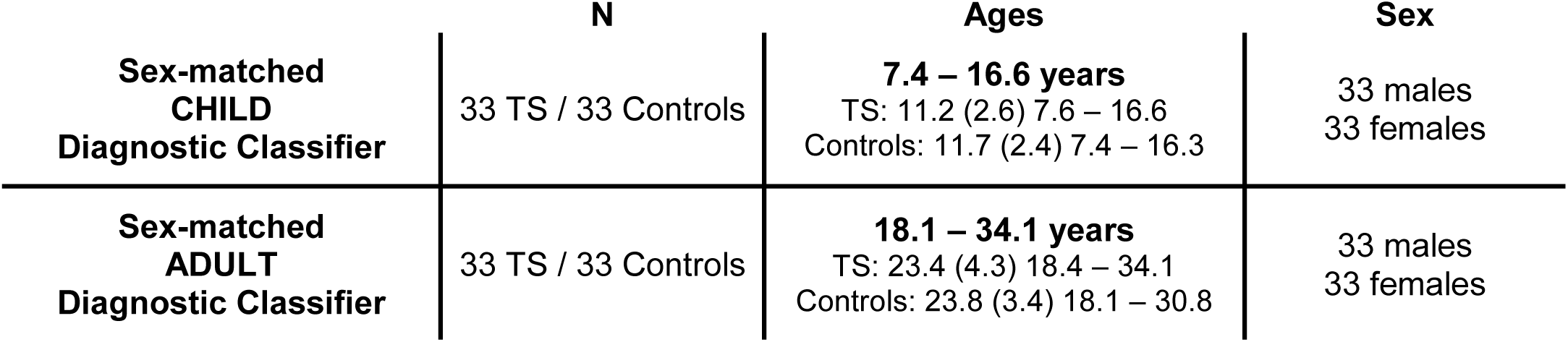
Training sets for sex-matched CHILD and ADULT diagnostic classifiers

|  | N | Ages | Sex |
| --- | --- | --- | --- |
| **Sex-matched**<br>**CHILD**<br>**Diagnostic Classifier** | 33 TS / 33 Controls | **7.4 – 16.6 years**<br>TS: 11.2 (2.6) 7.6 – 16.6<br>Controls: 11.7 (2.4) 7.4 – 16.3 | 33 males<br>33 females |
| **Sex-matched**<br>**ADULT**<br>**Diagnostic Classifier** | 33 TS / 33 Controls | **18.1 – 34.1 years**<br>TS: 23.4 (4.3) 18.4 – 34.1<br>Controls: 23.8 (3.4) 18.1 – 30.8 | 33 males<br>33 females |

The sex-matched CHILD diagnostic classifier (7.4-16.6 years; Table S5) was 64% accurate with LOOCV, akin to the YOUTH diagnostic classifier (also 64%). The sex-matched ADULT diagnostic classifier (18-31 years; Table S5) was 88% accurate with LOOCV. All diagnostic classifiers were accurate significantly above chance, which is 50% (sex-matched CHILD: *p* = 0.01; sex-matched ADULT: *p* < 0.001). However, these sex-matched classifiers still did could not accurately distinguish TS from controls as well in the other age group. Specifically, the sex-matched CHILD diagnostic classifier was 56% accurate for classifying diagnosis in the sex-matched adults, and the sex-matched ADULT diagnostic classifier was 59% accurate for classifying diagnosis in the sex-matched children. These classifiers performed significantly worse in the other age groups (see Figure S1). Thus, the poor generalizability observed in Figure 2 is likely due to age-related differences rather than sex differences.

**Figure S1.**
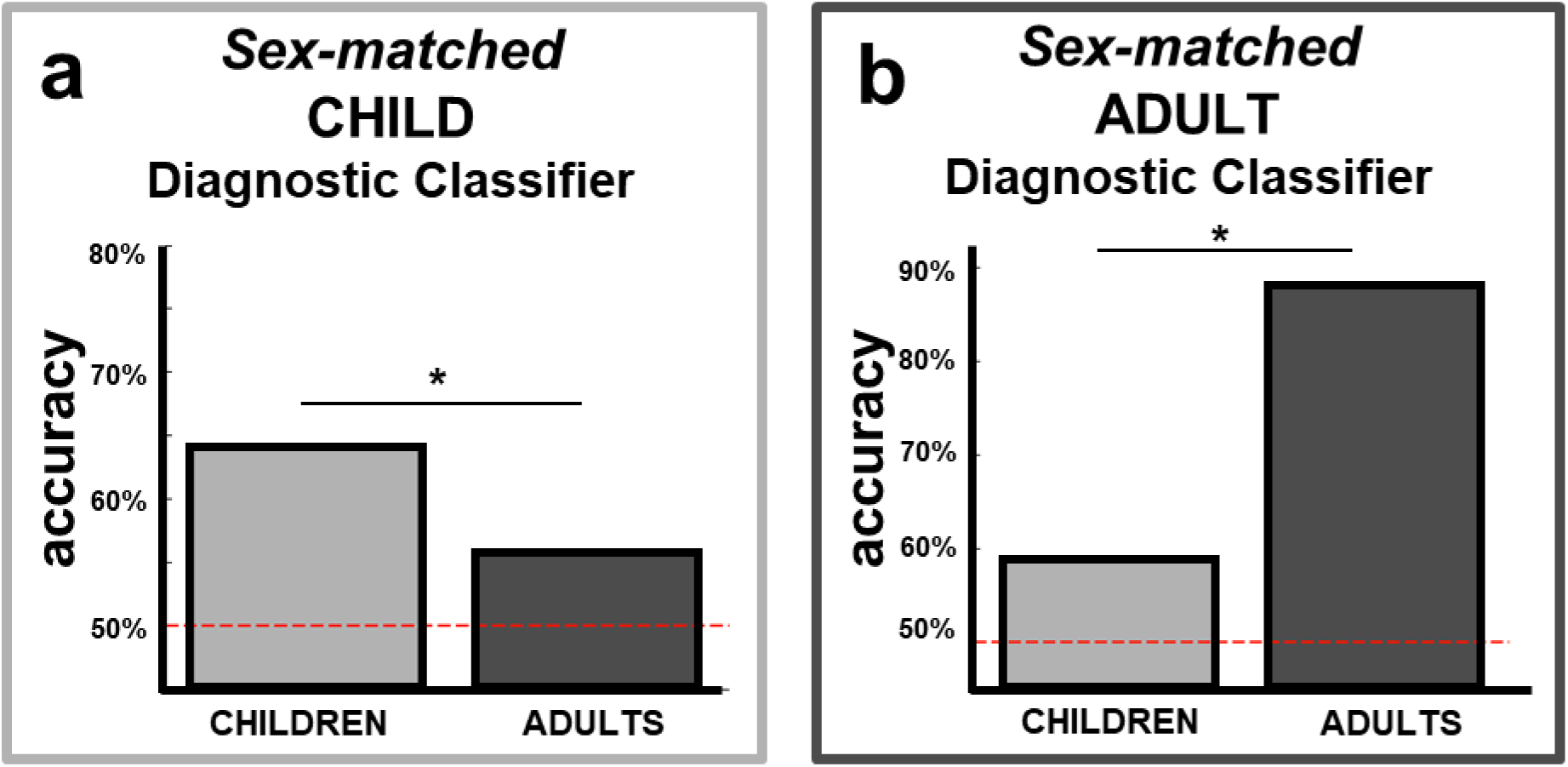
Functional connections that best distinguish TS from controls were age-specific, even when age groups were matched on sex. **a**.) Performance of the sex-matched CHILD diagnostic classifier to classify adults was significantly less accurate than performance in children (*p* = 0.01). **b.)** Performance of the sex-matched ADULT diagnostic classifier to classify children was significantly less accurate than performance in adults (*p* < 0.001).

### 2.2 ALL-AGES Diagnostic Classifier

To specifically target age-invariant differences in functional connectivity, we used SVM to build an ALL-AGES diagnostic classifier trained to distinguish TS from controls across the age range of our subjects (ages 7.4-34.2 years; Table S5). If some of these age-invariant differences are utilized by the CHILD or ADULT diagnostic classifiers, the CHILD and ADULT classifiers will generalize across age groups at least as well as the ALL-AGES diagnostic classifier. We used a binomial significance test to determine whether the performance of the CHILD or ADULT diagnostic classifiers significantly differed from the ALL-AGES diagnostic classifier (Supplemental Methods, *Binomial Significance Test*).

**Table S5.**
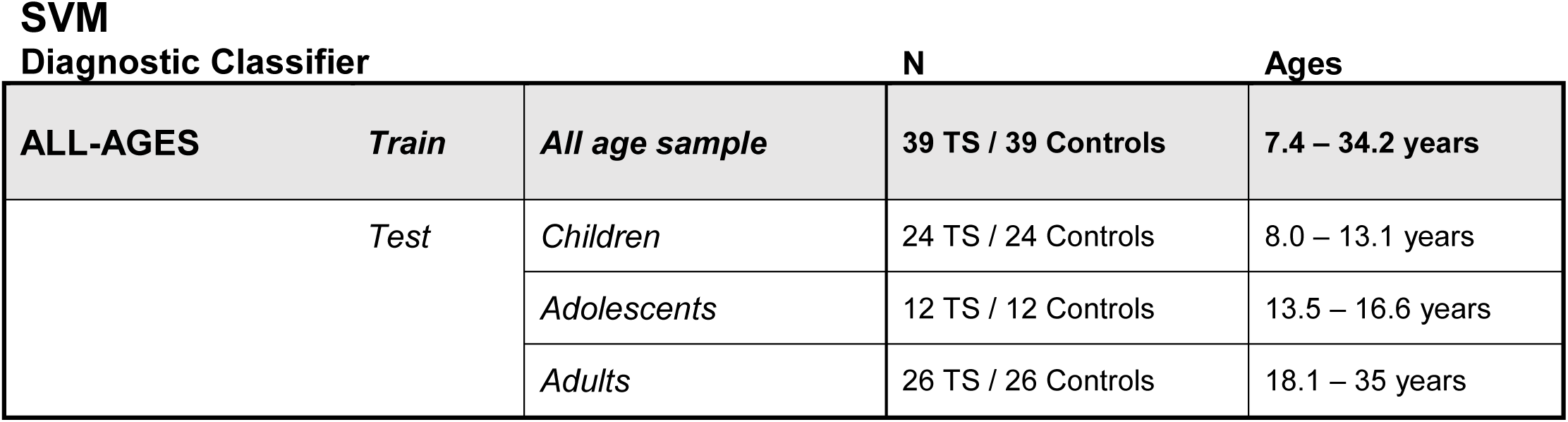
Training and testing sets used for the ALL-AGES diagnostic classifier

The ALL-AGES diagnostic classifier, which included children, adolescents, and adults (7-31 years; Table S5) was 60% accurate, marginally significant (*p* = 0.05). As the CHILD and ADULT diagnostic classifiers significantly outperformed the ALL-AGES diagnostic classifier (*p* = 0.015), the patterns of functional connections that best distinguished TS from controls in children and adults separately included age-specific differences.

**Figure S2.**
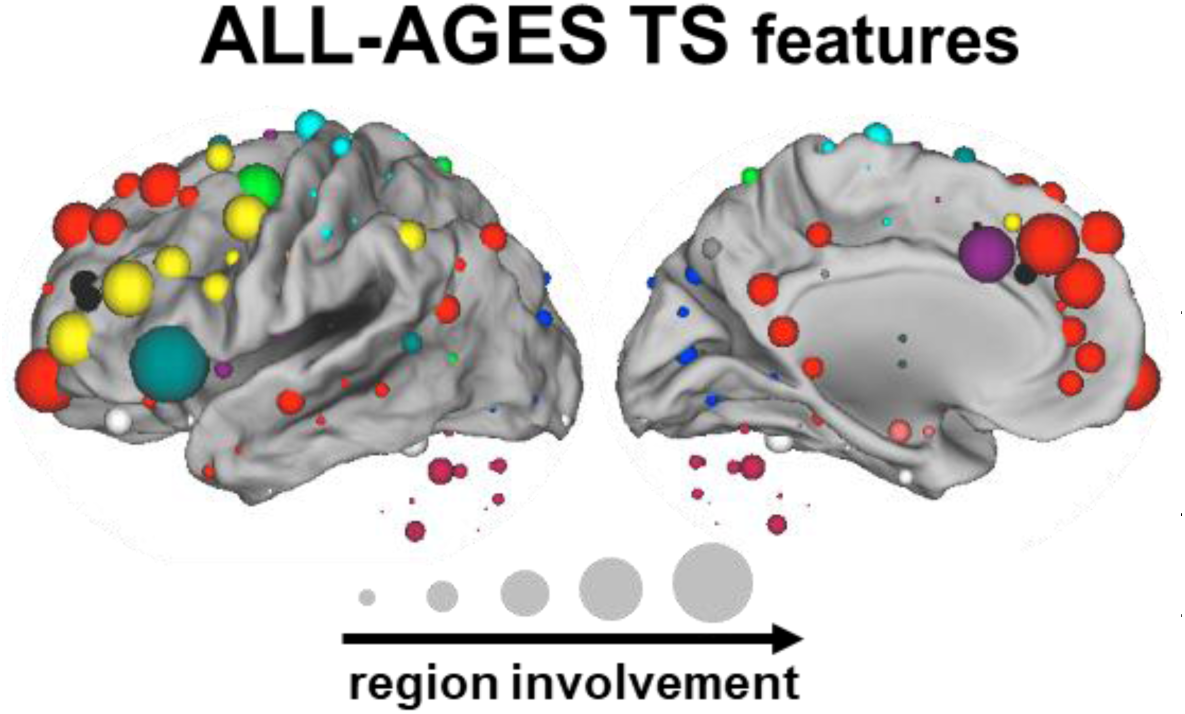
Functional connections that best distinguished TS from controls across children, adolescents, and adults. Regions are shown from the top 1000 weighted functional connections used to distinguish TS from controls in the ALL-AGES diagnostic classifier. The size of each sphere represents region involvement (i.e., number of functional connections from the feature set involving a particular region). Region colors indicate the network to which that region belongs.

We extracted the top 1000 (out of 44,850) functional connections most strongly weighted in the ALL-AGES diagnostic classifier. Regions involved in these functional connections were distributed among many processing, control, and default mode networks (Figure S2).

### 2.3 CHILD TS and ADULT TS Features

The top 1000 (out of 44,850) functional connections most strongly weighted in the CHILD or ADULT diagnostic classifiers were extracted. The top weighted functional connections are displayed in Figure S3 and show that these functional connections were within and between many different functional networks. Only 33 (3%) of the top 1000 functional connections overlapped between the CHILD and ADULT diagnostic classifiers. Top functional connections appear to be organized loosely by “blocks” of functional connections, either within a single network or between a pair of networks. Some blocks appear more heavily weighted in the ADULT diagnostic classifier (e.g., THAL-VIS or SM_body_-VIS) and other blocks appear to have different portions more heavily weighted by the CHILD and ADULT diagnostic classifiers (e.g., DMN). The different patterns of functional connections involved in the CHILD and ADULT diagnostic classifiers provide further evidence that TS differs between children and adults.

**Figure S3.**
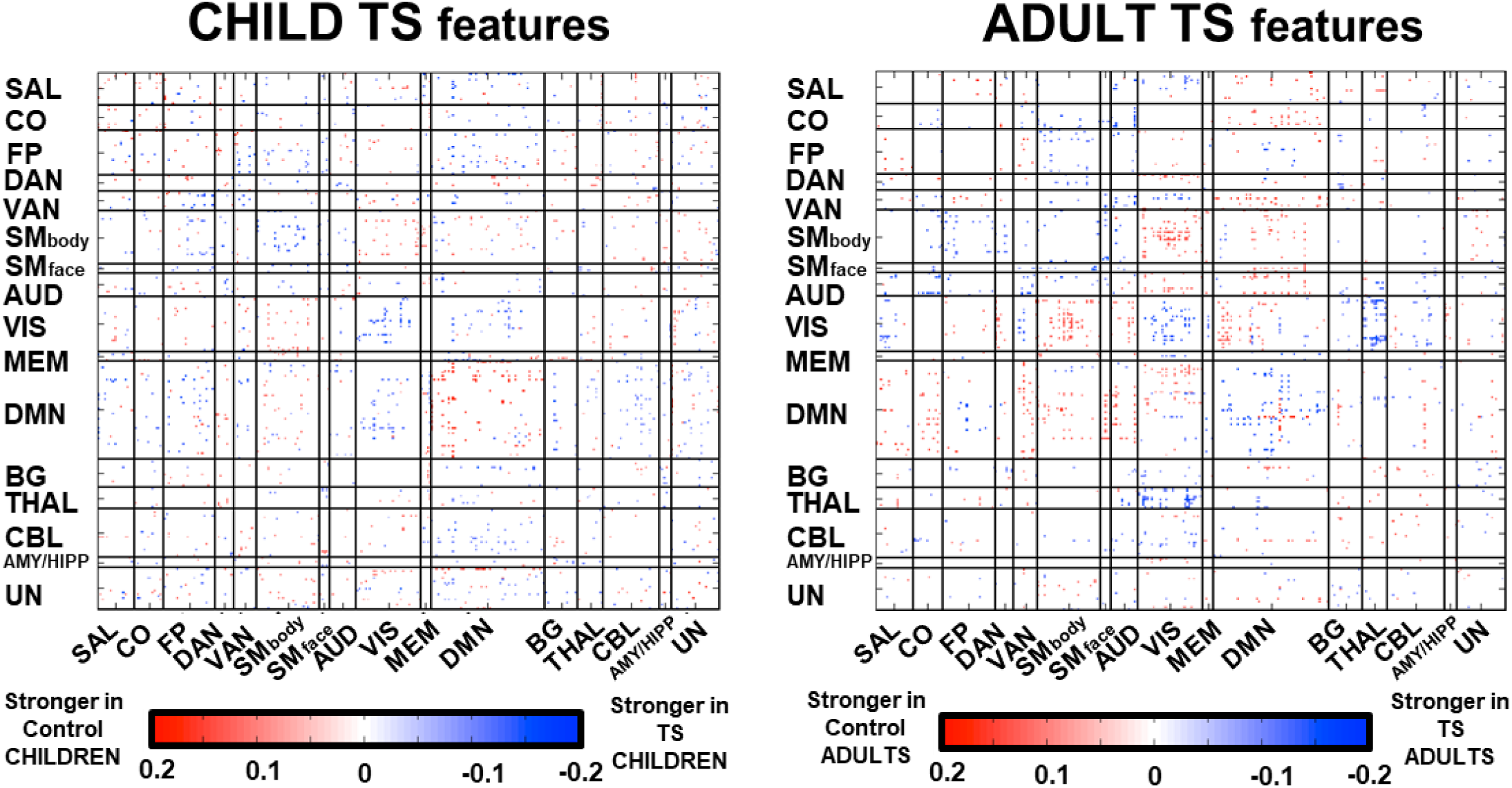
Functional connections selected as CHILD TS and ADULT TS features. Functional connections are shown from the top 1000 weighted functional connections used to distinguish TS from controls in the CHILD diagnostic classifier (*left*) and in the ADULT diagnostic classifier (*right).* Functional connections are shown between regions that are sorted by functional network and then from left to right. The average difference between TS and controls is depicted for each connection in both children (*left*) and adults (*right*).

### 2.4 Age Prediction with Random Features

In Nielsen et al. 2018, we found that many sets of functional connections, even randomly selected, can be used to predict the age of typically developing individuals, using SVR. In fact, in some cases, randomly selected features outperformed features selected based on *a priori* hypotheses. To evaluate whether the CHILD TS and ADULT TS features (i.e., those features most heavily weighted in each of these diagnostic classifiers) carry sufficient information related to age, we wanted to ensure that a matched number of randomly selected features did not outperform the disorder-related features. One hundred sets of 1000 randomly selected functional connections were used to build developmental models to predict age in the controls using SVR. Leave-one-out crossvalidation was used to assess performance of each developmental model. Figure S4 shows the amount of age-related variance explained by the predicted ages in controls from each set of randomly selected features, and how this compares to the CHILD TS and ADULT TS features. Randomly selected features did not outperform the CHILD TS and ADULT TS features and, thus, these disorder-related features carry sufficient information related to age.

**Figure S4.**
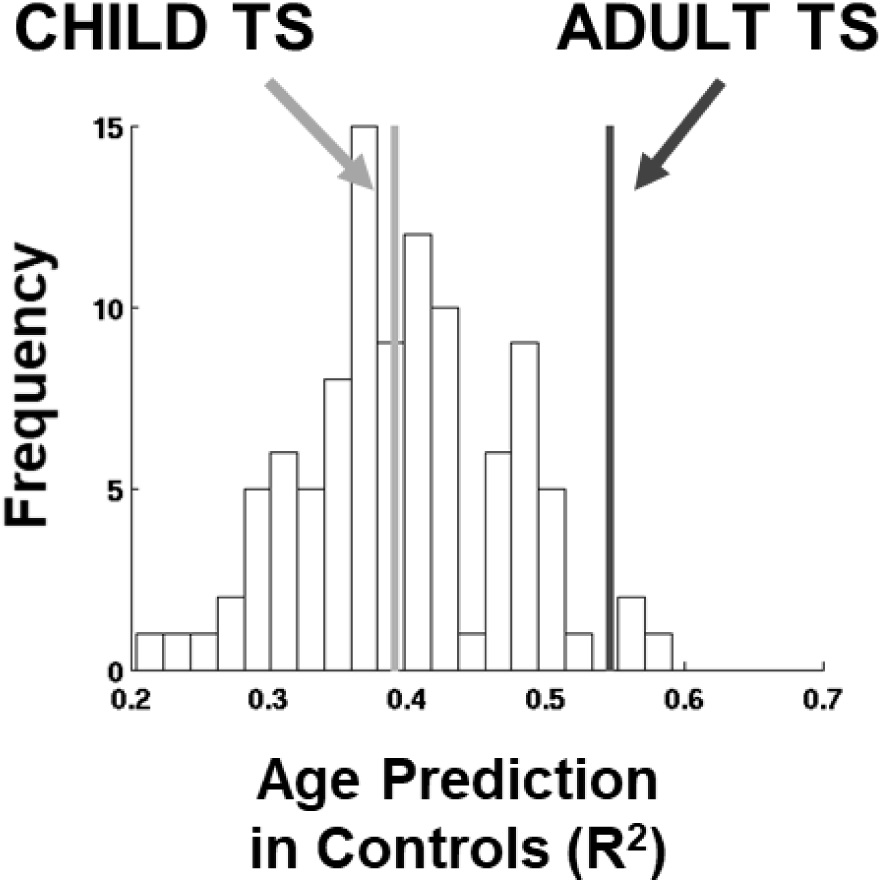
CHILD TS and ADULT TS features have sufficient information to predict age in controls. Variance explained by age in 100 developmental models built from 1000 randomly selected functional connections using SVR is displayed. The performance of the developmental models built from the CHILD TS and ADULT TS features is on par with those built from randomly selection connections.

### 2.5 Intact Development in TS

To further explore typical development of functional connectivity in TS, we used SVR trained on other feature sets in the controls to predict age in the TS participants. First, we included all possible features (all functional connections among the 300 regions across the whole brain), and as in Nielsen et al. 2018, we found that whole-brain functional connectivity contained age-related patterns that could be used to predict age well in controls (*r* = 0.73, *R*^2^ = 0.54, *p* < 0.001). These age-related patterns were maintained in TS, as the model also predicted age well in the TS group (*r* = 0.71, *R*^2^ = 0.50, *p* < 0.001). Second, we included the features that change the most in typical development by selecting the top 1000 features from a developmental classifier trained on 39 control children and 39 control adults (Table 2, controls used to train the CHILD and ADULT diagnostic classifiers) using SVM. These functional connections predicted age well in controls (*r* = 0.74, *R*^2^ = 0.56, *p* < 0.001) and in TS (*r* = 0.62, *R*^2^ = 0.38, *p* < 0.001), indicating that the age-related patterns in these developmentally relevant functional connections appear to be largely intact in TS.

Finally, we used SVR to predict age from the functional connections that differ most in the ALL-AGES diagnostic classifier, i.e., the top-weighted 1000 functional connections from the ALL-AGES diagnostic classifier. We found that these functional connections predicted age well in controls (*r* = 0.57, *R*^2^ = 0.32, *p* < 0.001) and in TS (*r* = 0.57, *R*^2^ = 0.32, *p* < 0.001), indicating that the developmental differences in the connections exhibiting age-invariant TS effects were largely intact in TS.

### 2.6 False Age Prediction

Figure 4 in the main text indicates that the predicted ages of TS individuals differ from the predicted ages of controls. This offset between the actual age and predicted age of TS individuals might arise if (1) TS individuals lack the age-related patterns of functional connectivity differences identified by the developmental model or (2) TS individuals have functional connectivity that exhibits accelerated or delayed maturation. To sort out these possibilities, we used SVR trained on the CHILD TS or ADULT TS features to build developmental models (n=100) to predict age in the controls using false, permuted age labels. Then, we tested how these false developmental models predicted age in the TS participants. As expected, false developmental models did not predict the age of TS individuals well (CHILD TS: average R^2^ = 0.021; ADULT TS: average R^2^ = 0.044). Rather, the predicted ages fell near the mean age of the training set, indicating a failure of the model (Figure S5, *grey).* Further, these predicted ages differed from the predicted ages of TS individuals when the real (not false) ages of the controls were used to the build the model (see main text), indicating that the functional connections that differ in TS reflect shifted development.

**Figure S5.**
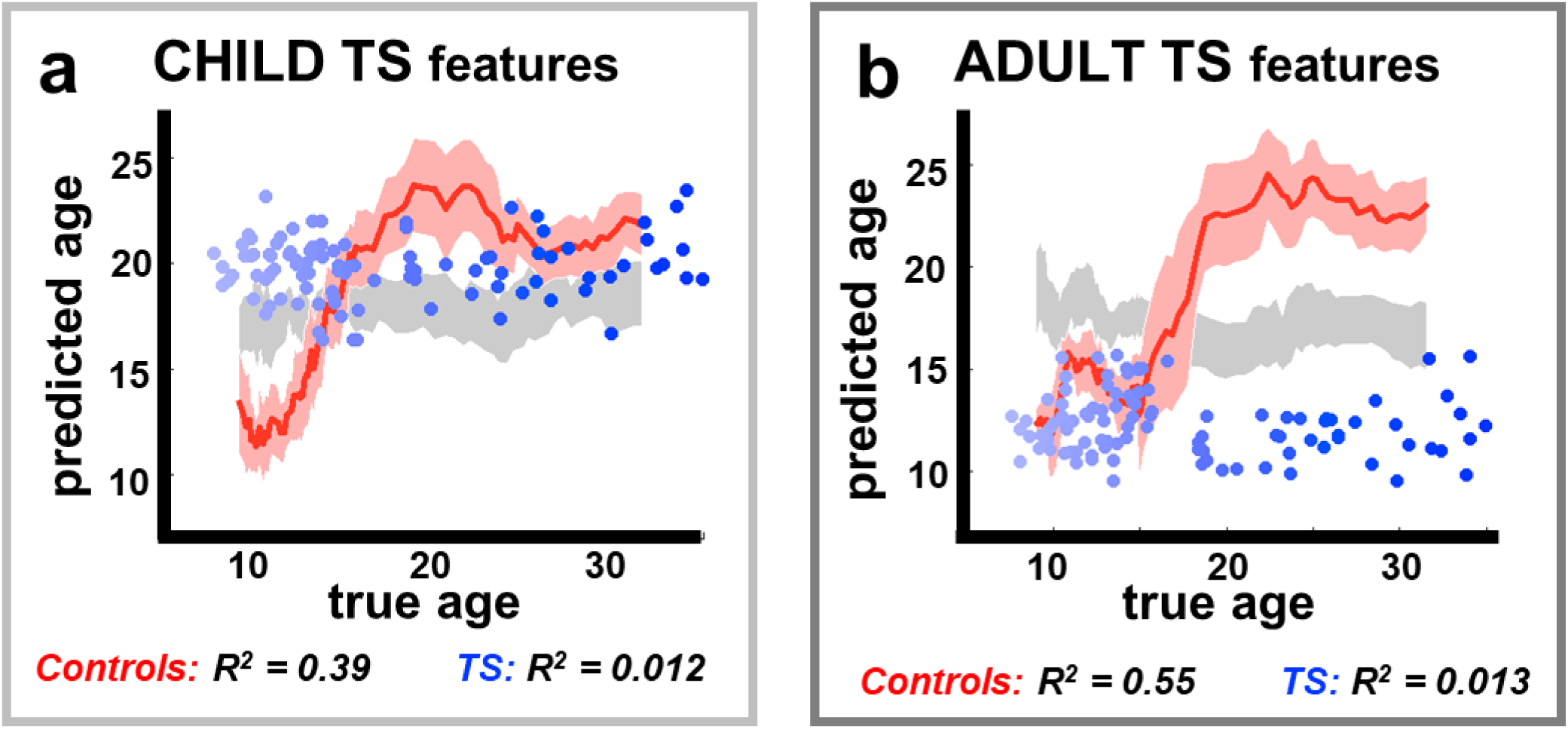
Predicted ages with CHILD TS and ADULT TS features reflect shifted rather than absent development. **a.**) The developmental model built using the CHILD TS features was able to predict age well in the control sample (*red*) but not in the TS sample (*blue*). The false developmental models using CHILD TS features did not predict age well in the TS sample (*grey*). Predicted ages of children with TS were older than the predicted ages of age-matched controls and older than their predicted ages form the false developmental models, indicating acceleration maturation in the CHILD TS features. **b.**) The developmental model built using the ADULT TS features was able to predict age well in the control sample (*red*) but not in the TS sample (*blue*). The false developmental models using the ADULT TS features did not predict age well in the TS sample (*grey*). Predicted ages of adults with TS were younger than the predicted ages of age-matched controls and younger than their predicted ages from the false developmental models, indicating delayed or incomplete maturation in the ADULT TS features.

